# Reconstruction of sound driven, actively amplified and spontaneous motions within the tree cricket auditory organ

**DOI:** 10.1101/2021.11.14.468538

**Authors:** Natasha Mhatre, James B. Dewey, Patricia M. Quiñones, Andrew Mason, Brian E. Applegate, John S. Oghalai

**Affiliations:** Department of Biology, Western University, London, Ontario. Canada; Brain and Mind Institute, Western University, London, Ontario. Canada; Department of Biological Sciences, University of Toronto at Scarborough, Scarborough, Ontario, Canada; Caruso Department of Otolaryngology – Head and Neck Surgery, University of Southern California, Los Angeles, California, USA

**Keywords:** insect hearing, tree crickets, active amplification, optical coherence tomography, ciliary beating

## Abstract

Hearing consists of a delicate chain of events. Sound is first captured by an eardrum or similar organ which is set into vibrations, these vibrations must then be transmitted to sensory cells in a manner that opens mechanosensory channels generating an electrical signal. Studying this process is challenging. Auditory vibrations are in the nano- to picometer-scale and occur at fast temporal scales of milli to microseconds. Finally, most of this process occurs within the body of the animal where it is inaccessible to conventional measurement techniques. For instance, even in crickets, a century-old auditory model system, it is unclear how sound evoked vibrations are transmitted to sensory neurons. Here, we use optical coherence tomography (OCT) to measure how vibrations travel within the auditory organ of the western tree cricket (*Oecanthus californicus*). We also measure the reversal of this process as mechanosensory cells generate spontaneous oscillations and amplify sound-evoked vibrations. Most importantly, we found that while the mechanosensory neurons were not attached to the peripheral sound collecting structures, they were mechanically well-coupled through acoustic trachea. Thus, the acoustic trachea are not merely conduits for sound but also perform a mechanical function. Our results generate several insights into the similarities between insect and vertebrate hearing, and into the evolutionary history of auditory amplification.

## Introduction

The rich soundscape of the summer nights is familiar in almost all parts of the world. In this chorus, insects dominate the sounds humans can perceive. The predominant sound is that of cricket and katydid males trying to attract females. Among the listeners, the female crickets and katydids using these sounds to evaluate and track down mates aren’t alone (Pollack, 2000). At the same time, eavesdropping flies will use the very same sounds to find the singing males and parasitize them (Mason et al., 2001). In yet another layer of complexity, there are sounds that humans cannot hear, made by predatory bats hunting for their insect prey. While we cannot, many insects can hear the bats and the bat’s echolocation calls trigger quick escape responses (Conner and Corcoran, 2012).

Thus, hearing is crucial to both reproduction and survival in insects. This range of evolutionary pressures has led to the independent evolution of a diversity of tympanal or eardrum-based ears in at least 17 different insect lineages (Hoy and Robert, 1996), in addition to antennal ears used by some Dipterans (Nadrowski et al., 2011). Indeed, insect ears appear on a variety of body parts and have distinct anatomies and mechanical specializations (Windmill and Jackson, 2016). All of them, regardless of their anatomical location, rely on a peripheral organ that converts sound into a mechanical motion, which is then transmitted to mechanosensory neurons (Yack, 2004; Yager, 1999). These neurons then transduce this motion into an electrical signal which can be transmitted to other parts of the nervous system and processed to determine the animal’s response to the perceived sound (Hedwig, 2014, 2013).

One of the longest and best-studied insect model systems in the field of naturalistic acoustic communication are the true crickets or the Gryllidae (Pollack and Noji, 2017; Regen, 1913). A great deal is known about the biophysics of cricket ears (Michelsen and Larsen, 1978), the nervous systems that process auditory information (Hedwig, 2014), the behavior of these insects in the field (Mhatre and Balakrishnan, 2008; Nandi et al., 2019; Rodríguez-Muñoz et al., 2010) and even about the rapid and interesting evolutionary pathways taken by their communication systems(Pascoal et al., 2014; Tinghitella et al., 2021). A significant gap in our knowledge of true crickets, however, is that we do not understand the first step in the chain of hearing. In crickets, auditory mechanosensory neurons are not attached to the eardrum, and we do not know how the motion of this peripheral sound-collecting organ, the eardrum or tympanum, is delivered to the neurons that transduce sound. Indeed, this is important because crickets, and their sister group the tettigoniids, are known to possess two crucial features of vertebrate hearing, active amplification (Mhatre and Robert, 2013; Morley and Mason, 2015) and tonotopy (Montealegre-Z. et al., 2012; Oldfield et al., 1986; Palghat Udayashankar et al., 2012). Thus, understanding the nature of force transmission in cricket auditory systems will enable us to fully exploit their potential as more broadly applicable model systems for hearing.

The auditory organ of crickets is on their forelegs and the tympana are conveniently exposed and amenable to conventional vibrometry. The mechanosensory neurons, however, are internal structures, and direct optical access is unavailable. To date, the vibrational behaviour of mechanosensory neurons has largely been inferred based on the motion of the external sound collecting organs (Mhatre, 2015). However, with the advent of optical coherence tomography (OCT) based vibrometry, it is now possible to measure the motion of internal structures at the spatial and temporal resolutions relevant to auditory systems (Dewey et al., 2019; Lee et al., 2015). OCT can resolve internal structures that are in the micrometer size range, and its ability to detect vibration is significantly more sensitive, lying in the pico- to nano-meter range and at very high speeds (∼200 kHz). Here we use volumetric optical coherence tomography vibrometry or VOCTV (Lee et al., 2015) to study insect hearing for the first time. Using VOCTV, we simultaneously measure the motion of the external and internal structures of the tympanal organ of a member of the Gryllidae, the western tree cricket (*Oecanthus californicus*).

We chose a subgroup of true crickets called the tree crickets (family: Gryllidae, subfamily: Oecanthinae) as a model system for this study mainly because they possess active auditory amplification (Mhatre et al., 2016; Mhatre and Robert, 2013). Indeed, they are among the few insects confirmed to possess active amplification; their tympana show spontaneous oscillations (Mhatre and Robert, 2018, 2013; Morley and Mason, 2015), and these oscillations dissipate power, a clear sign of true active amplification (Mhatre, 2015; Mhatre and Robert, 2018). It is this feature that makes them a promising model system for a study of auditory organ mechanics; in tree crickets we can study in a minimally invasive way, not only how sonic mechanical energy is delivered to mechanosensory neurons, but also measure the reversal of the process during active amplification; thus we can study both forward and reverse force transmission. Here, using VOCTV, first we generate a 3D reconstruction of the western tree cricket auditory organ. Based on this 3D model and on vibrational motion at conspecific call frequency, we identify the structures to which mechanosensory neurons are attached. We then use measurements from these attachment points, from two perpendicular views to reconstruct the 2D motion of the auditory organ during sound driven and mechanosensory neuron driven vibration.

## Results

### 3D structure

Using the reflectivity data we get from the VOCTV system, we can reconstruct the 3D structure of the auditory organ of the western tree cricket (*O. californicus)* (Fig.1). Such a reconstruction enables a comparison to the organ morphology known from the auditory organs of other true crickets (Family: Gryllidae) (Michel, 1974; Oldfield et al., 1986; Schneider et al., 2017). In subsequent measurements, the 3D reconstruction enables us to select the correct measurement planes and points within the organ in order to reconstruct its motion.

**Figure 1.**
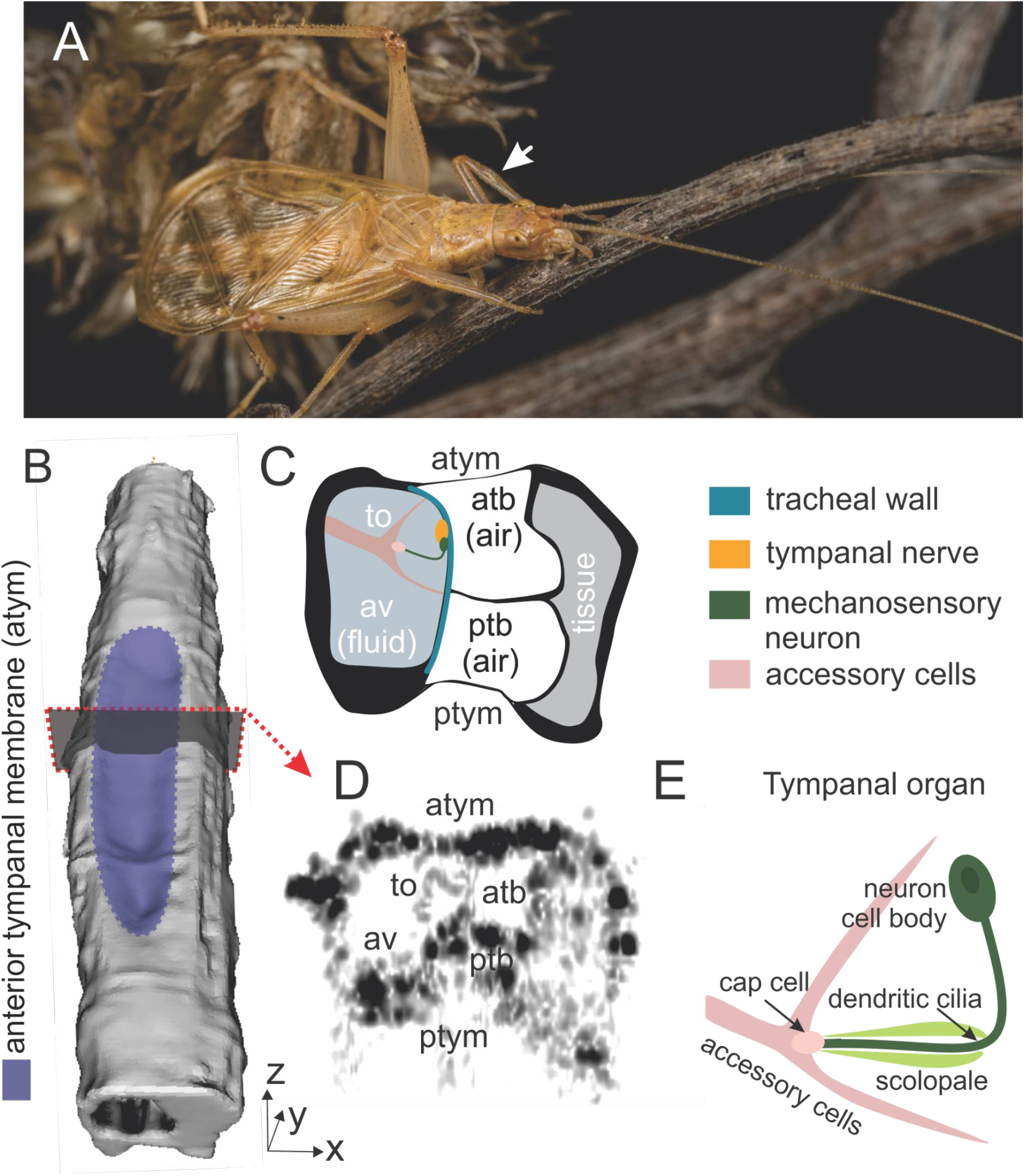
OCT captures crucial morphological features within the auditory system of the western tree cricket (Oecanthus californicus). (A) The auditory organ in tree crickets is positioned on their foreleg and indicated by the arrow in this image of a male western tree cricket (image courtesy Alice Abela). Externally, each tree cricket ear includes two large tympanal membranes, the anterior tympanal membrane (atym) which is exposed here, and a posterior tympanal membrane (ptym) which is positioned on the posterior side of the foreleg. (B) A 3D reconstruction of the auditory organ on the foreleg (arrow in panel A) was made based on OCT reflectivity data. (D) A section of this data through the leg shows the positions of the anterior tympanal membrane and also enables us to locate a few crucial internal structures known from other gryllid ears. These structures are marked in (C): the auditory vesicle (av), a hollow fluid filled cavity which houses the tympanal organ (to), the air-filled anterior (atb) and the posterior tracheal branch (ptb) of the acoustic trachea. These are air filled hollow tubes that lie under the two tympanal membranes and create a low impedance air cavity that allows for tympanal motion. The cricket tympanal organ (to) contains mechanosensory neurons that transduce the motion of the tympanum into neural signals. Additionally, the organ includes an array of accessory cells that provide functional and structural support, attaching the organ to the tracheal and the auditory vesicle walls. The tympanal nerve, shown in yellow, carries the axons of the mechanosensory neurons to the CNS. The nerve lies within the auditory vesicle and runs along the external wall of the auditory trachea, which is shown in blue. The mechanosensory neurons, shown in green, emerge from this nerve and are connected via accessory cells to the internal cuticle of the auditory vesicle. We will use the 3 dimensional x-y-z convention established in this figure throughout this manuscript. (D) In this slice taken from the OCT reflectivity data, which was used to make the 3D reconstruction (B), we can observe some of these components of the auditory organ, specifically, the two tympanal membranes (atym, and ptym), the anterior and posterior branches of the acoustic trachea (atb, ptb) and, parts of the tympanal organ (to). Darker colours indicate higher reflectivity. The spatial resolution here (∼10 μm axial and lateral) does not reflect the vibrometric resolution which is much higher (in the nano to picometer range depending on the frequency). (E) A schematic showing the sectional substructure of the tympanal organ in more detail. The tympanal organ contains mechanosensory units known as scolopales, complex structures which contain mechanosensory neurons and supporting scolopidial cells. In the neuron, it is the dendritic cilium which is the site of auditory transduction and active amplification. These mechanosensory cilia are made of 9 microtubule pairs arranged in a wheel-like fashion, but there is no central pair. Motor molecules known as axonemal dyneins connect adjacent microtubule pairs. Typically, the activity of axonemal dyneins causes the rod-like microtubules which run through the length of the cilium to slide against each other, and cause the structure to bend and beat. In insect mechanosensory neurons, it is these axonemal dyneins that have been shown to be responsible for the motor function underlying active amplification in *Drosophila melanogaster*.

Typically, the gryllid ear is located on the tibia of the foreleg (Fig. 1A, B) and has two external eardrums known as tympanal membranes. These can be of different relative sizes depending on the sub-family (Schneider et al., 2017). In tree crickets, the anterior tympanal membrane (atym) which is on the anterior face of the leg, is of similar size to the posterior tympanal membrane (ptym) and has a similar frequency response (Mhatre et al., 2009). Inside the foreleg, there is a large diameter trachea known as the acoustic trachea, which serves as a secondary sound input to the organ (Michelsen and Larsen, 2008), and reduces the impedance to tympanal vibration at appropriate frequencies. This trachea splits into two branches, the anterior tracheal branch (atb) and the posterior tracheal branch (ptb) (Fig. 1C). These two branches are resolved with OCT (Fig. 1D). On the lateral side of the foreleg, opposed to the two tracheal branches, is the auditory vesicle (av, Fig. 1C) which is also clearly resolved in the OCT image (Fig. 1D).

The auditory vesicle houses the tympanal organ, which is responsible for auditory transduction. This organ contains bipolar mechanosensory neurons, which transduce sound driven motion into neural signals, and accessory cells which provide physiological and structural support (Fig. 1C-E). The neurons themselves possess a cell body, a ciliated dendrite and an axon. Transduction, as well as active amplification takes place in the ciliated dendrite in insects (Hehlert et al., 2020). On the other side of the bipolar neuron, the axon fuses with the tympanal nerve, which then carries these signals upstream. This nerve which lies inside the auditory vesicle, travels along the external wall of the two tracheal branches (Fig 1, 2). Thus, the tympanal organ is connected to the wall of the acoustic trachea by the tympanal nerve, and by the accessory cells to the internal wall of the auditory vesicle (Fig 1, 2). The organ stretches across the auditory vesicle, but does not make direct contact with either tympanal membrane in gryllid auditory organs (Michel, 1974; Oldfield et al., 1986; Schneider et al., 2017).

**Figure 2:**
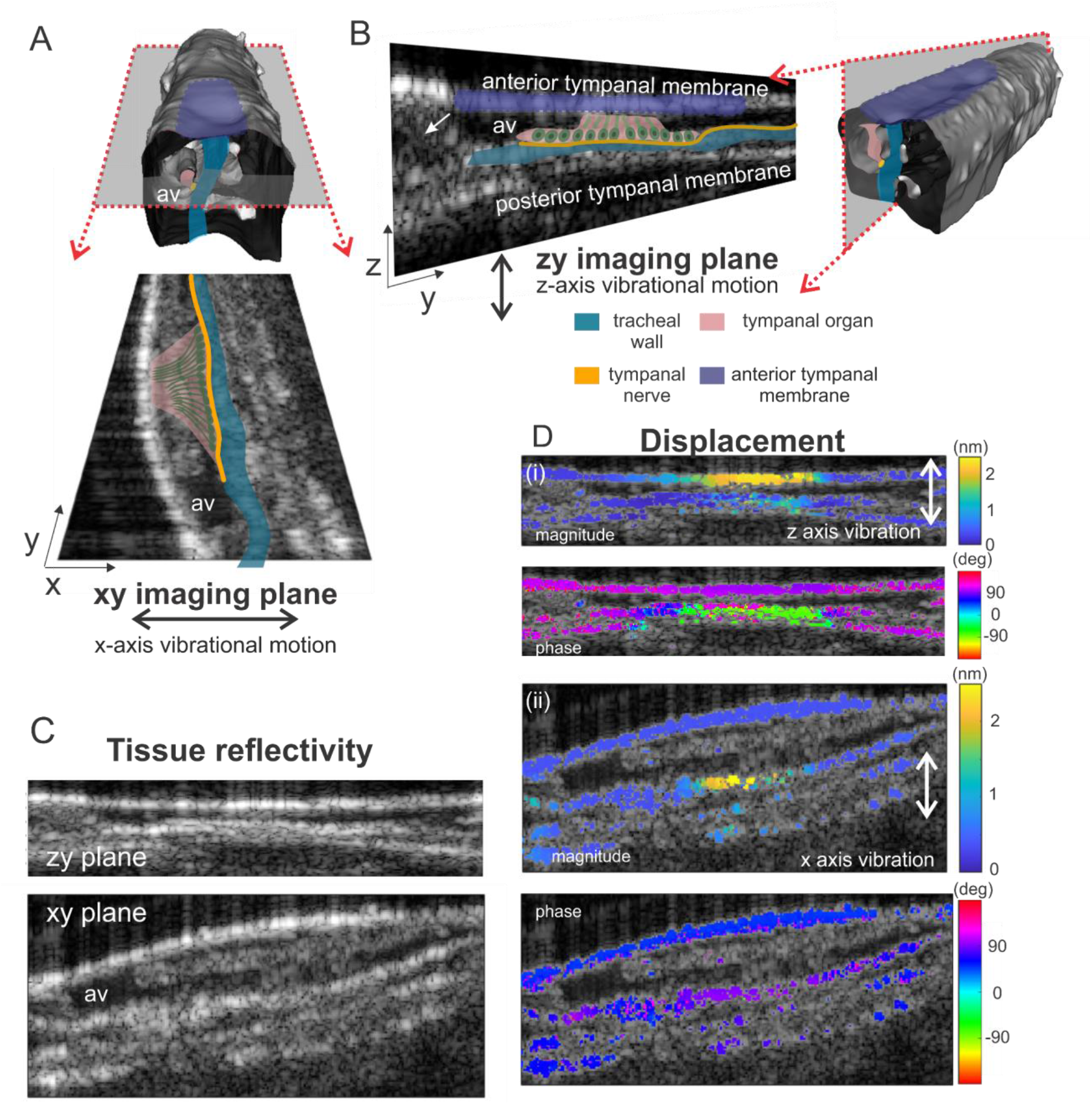
Measurement planes within tree cricket auditory organ simultaneously capture the motion of the tympanal membranes and the tympanal organ attachments. (A) From the 3D structure, we identified two important measurement planes within the auditory organ. An array of the mechanosensory neurons identified in the previous figure are known to ‘fan’ out from the tympanal nerve and lie approximately within a plane that is parallel to the anterior and posterior tympanal membranes. These neurons are connected via accessory cells to the internal cuticle of the auditory vesicle. This structure together is known as the tympanal organ. The tympanal organ has a complicated shape but the mechanosensory scolopale units within it lie largely within an x-y planar section. Measurement from an appropriately chosen x-y plane can identify lateral (x-axis) motion in the scolopales and hence of the dendritic cilia within it. (B) Measurements in this z-y plane allows us to measure the vertical (z-axis) motion of the anterior tympanal membrane and simultaneously from the internal tracheal wall where the scolopales are attached. (C) Both cuticle and trachea show excellent reflectivity (bright regions) and we are readily able to identify these internal structures as well as the air cavities (dark regions) within them. Softer tissues like the cell masses and fluids within the auditory vesicle (av) have lower reflectivity and appear greyer than the cuticle and trachea. (D) We only consider the vibrometric data from regions of high reflectivity, where we have sufficient signal to reliably estimate displacement. The scans depicted here were carried out at the amplified frequency at a stimulation level of 80 dB SPL, which corresponds to conspecific song frequency (3.25 kHz at 21°C). (i) In the z-y imaging plane, we see significant vertical displacement of the anterior tympanal membrane as well as the internal tracheal wall. We also observe that the internal tracheal wall where the tympanal organ is attached is shifted nearly 180° in phase in comparison to the external tympanal membrane. Thus, when stimulated by conspecific frequency the external tympanal membrane and the internal tympanal organ will move in opposite directions, maximizing vertical deformation. (ii) In the x-y imaging plane, we see significant lateral displacement only at the tracheal wall, and here the phase is more similar to that observed at the anterior tympanal membrane.

The walls of the acoustic tracheal branches and the auditory vesicle are quite reflective, and well resolved by OCT providing us with reliable and very low noise vibrometric measurements. The tympanal organ itself is more transparent and does not reflect the OCT laser strongly (Fig. 1D, Fig 2). However, since there is good signal from the internal attachment points to which the organ is connected, we can still infer and reconstruct the likely motion of the organ and the mechanosensory neurons within it.

### 2D vibrometric measurements

We identified two planes within the organ which were crucial to reconstructing the organ’s motion (Fig. 2). As mentioned before, the tympanal organ stretches across the auditory vesicle. In gryllids, the organ is thought to lie roughly within a plane parallel to the two tympanal membranes. Within this plane, a fan-like array of mechanosensory cells emerges from the tympanal nerve and connects to a smaller area on the inner wall of the auditory vesicle *via* accessory cells (Michel, 1974; Oldfield et al., 1986). We selected an x-y plane parallel to the two tympanal membranes, which intersected the tympanal organ’s attachment to the acoustic trachea (Fig. 2A). This determination was based both on the anatomy, and the observation that this plane captured the point to highest vibrational motion in the x direction at conspecific song frequency (Fig. 2A, D).

Previous data have shown that tree cricket tympanal membranes are quite stiff and both sound-driven and spontaneous motion is concentrated to a relatively small section of the anterior tympanal membrane (Mhatre et al., 2009; Morley and Mason, 2015). Hence, the second measurement plane was a z-y plane that ran through the anterior and posterior tympanal membranes, and through the tracheal wall near the tympanal organ (Fig. 2B). The plane was chosen to capture the positions at which z displacement was maximal on the anterior tympanal membrane and the internal tracheal wall at conspecific frequency (Fig. 2B, C).

The data shown here were collected in response to sound stimulation close to conspecific call song frequency at 80 dB sound pressure level (SPL re 2e-5 Nm^-2^) (Fig. 2: using a 3.2 kHz tone). The patterns of external anterior membrane vibration in the western tree cricket are similar to those previously measured in other tree crickets. We also can make several new observations. First, the internal motions of the tympanal organ were concentrated within a small area, along the tracheal wall where the tympanal organ is connected. This part moves significantly in both x and z directions, indicating that this part undergoes complex motion in at least two dimensions (Fig. 2D).

Even more interestingly, we observed a ∼180° phase difference in the displacement of the external anterior tympanal membrane and internal tracheal wall in the z direction. Thus, when stimulated by conspecific frequency, the two membranes move in opposite directions, maximizing the vertical deformation applied to the tracheal wall. In the x direction, the displacement of the internal tracheal wall is closer in phase (∼90°) but not identical to that of the external tympanal membrane in the z direction. A more complex geometric approach will be needed to reconstruct the motion of the organ based both on these phase differences and the relative amplitudes of motion. In order to tackle this reconstruction, we will first consider detailed single point frequency response measurements averaged across a larger number of animals.

### Single point vibrometric measurements

Locations that vibrated maximally were chosen for displacement measurements. Displacements were measured in response to sound stimulation across frequencies and sound pressure levels. We measured motion in the z direction from the external anterior tympanal membrane (Fig. 3A) and the internal tracheal wall (Fig. 3B), and in the x direction from the tracheal wall (Fig. 3C). Like other tree crickets (Mhatre and Robert, 2013; Morley and Mason, 2015), the western tree cricket’s anterior tympanal membrane showed frequency-selective and stimulus amplitude-dependent displacement (Fig. 3 A). Measurements from the tracheal wall also showed similar non-linear mechanics in both x and y directions (Fig. 3 B, C).

**Figure 3.**
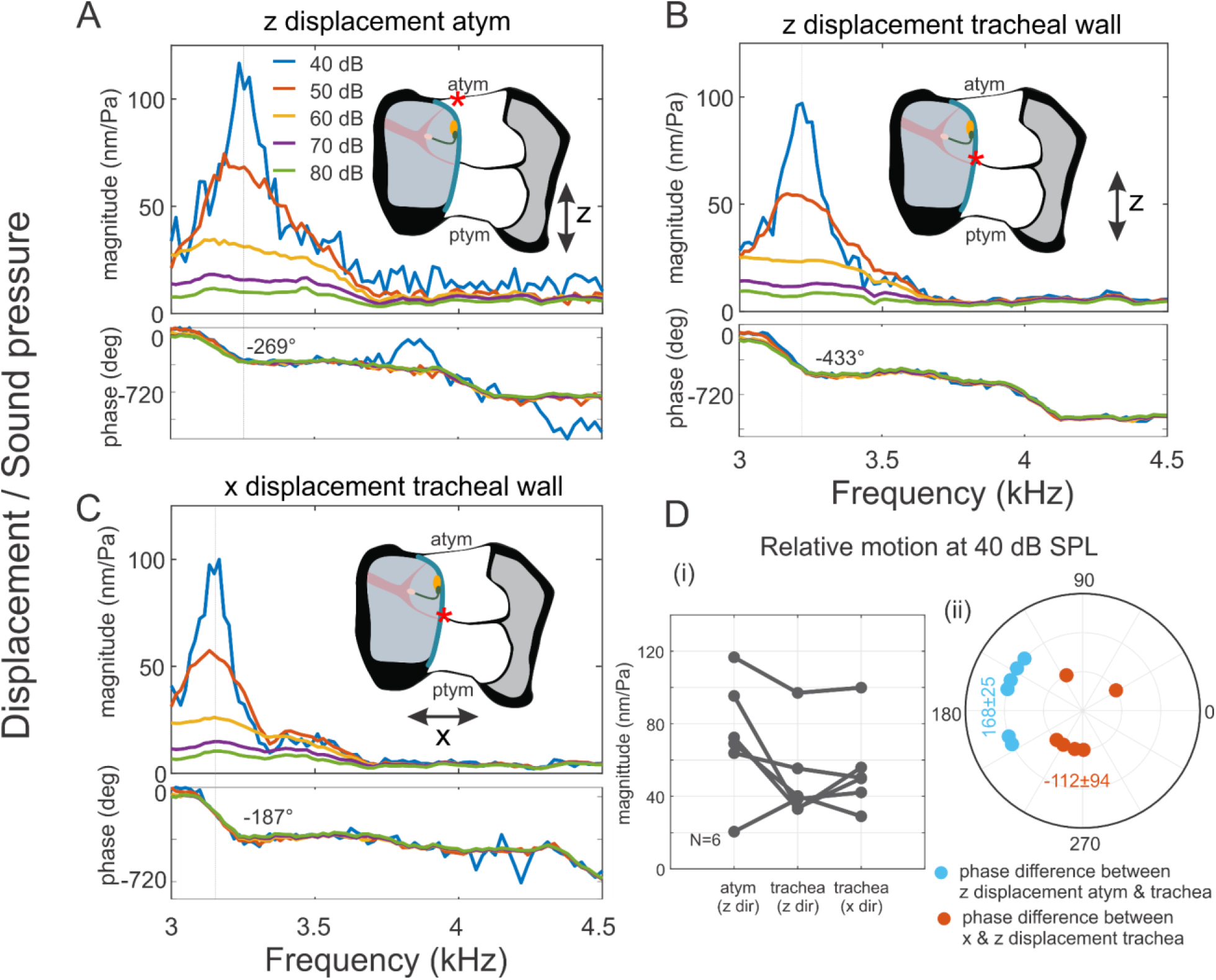
The frequency response of the tree cricket auditory organ shows compressive non-linearity. Using the 2D scans, we selected single points of maximal displacement on the (A) external anterior tympanal membrane, and (B, C) the internal tracheal wall (depicted by *). The sensitivity or displacement relative to sound pressure level was measured at these points at different stimulation levels, along the (A, B) z and (C) the x axis. We observe compressive non-linearity and level dependent tuning in all three measurements including at the attachment point of the tympanal organ. At 40 dB SPL, a large proportion of the force is produced by the mechanosensory neurons and in the individual depicted in A, B and C, the displacements are similar for all measurements. (D) If we compare the (i) peak sensitivity magnitude across individuals, we find that the external and internal magnitudes are similar in the z direction (73.0±32 nm/Pa vs 49.6±24.5 nm/Pa respectively) and the magnitudes of the internal membrane in the x and z direction are also similar (54.5±24.1 nm/Pa vs 49.6±24.5 nm/Pa respectively). (ii) The phase difference between the z displacement of the external anterior tympanal membrane and the internal tracheal attachment point is 168° ± 25° (N=6), and the phase difference between the x and z direction displacements of the internal tracheal attachment point is 138°± 94° (N=6) with two strong outliers.

In order to test the effect of stimulus SPL and of direction and position of measurement, we carried out a repeated measures ANOVA with a linear mixed effects model (lme4 package, RStudio, RStudio 2021.09.0 build 351, (Bates et al., 2015)) where both stimulus SPL and the three measurements (z direction displacement of the external membrane, the x and z direction displacement of the internal tracheal wall) were treated as independent variables and the independent variable was sensitivity or displacement per unit sound pressure level. We found that only stimulus SPL had a significant effect on the sensitivity (F(1,25)=84.77, P<0.001, N=6 individuals) and there was no effect of measurement direction or position (F(2,25)=1.85, P<0.177, N=6 individuals) and no interaction effect (F(2,25)=1.38,P=0.27, N=6 individuals).

In all three measurements, at low stimulation levels (40 dB SPL), we observed a sharp peak in sensitivity (3.48 ± 0.18 kHz, N=6; Fig. 3) near the conspecific song frequency, i.e. ∼3.25 kHz at 21°C (Walker, 1962). Thus the actively amplified frequency in *O. californicus*, as in other tree crickets, corresponds to the mating call of the species (Mhatre et al., 2016; Mhatre and Robert, 2013; Morley and Mason, 2015).

As observed in other insect and mammalian actively amplified auditory systems, when stimulus SPL is increased, this peak in the sensitivity grows broader and flattens (Fig. 3). In this individual, at the two highest stimulation levels 70 and 80 dB SPL, the sensitivity of the anterior tympanal membrane becomes nearly linear, and the sensitivity observed is low, ∼10 nm/Pa (Fig. 3A). At the lowest sound pressure level tested, 40 dB SPL (2 mPa), the sensitivity at the same position in this individual was much higher ∼116 nm/Pa. This pattern of higher sensitivity at low sound stimulation levels holds across animals (40 dB SPL: 73.0 ± 32 nm/Pa, 80 dB SPL: 9.48 ± 4.2 nm/Pa). It also suggests that at 40dB SPL, 85.53 ± 4.2 % (N=6) of the membrane’s displacement in the z direction is produced by the amplification mechanism rather than by the sound.

While the active amplification mechanism increased sensitivity at low sound levels compared to high levels, relative displacement levels within the organ did not change across stimulation levels as tested by the repeated measures ANOVA. At 40 dB SPL, where displacements within the auditory organ were dominated by the amplification mechanism, we found that the displacement of the anterior tympanal membrane in z direction, and of the tracheal wall in x and z direction were not significantly different (Fig. 3D). Thus, when we compared the sensitivity between the external anterior tympanal membrane and the internal tracheal wall in the z direction, they were not significantly different (Fig.3 D) and similarly, the displacement of the internal tracheal wall in the x and z direction were also not significantly different (Fig. 3D).

If measured at the same peak frequency, this pattern was also maintained at higher stimulation levels (80 dB SPL), where forces from sound predominated. The sensitivity of the external anterior tympanal membrane in the z direction was significantly lower 9.48 ± 4.23 nm/Pa but that of the internal tracheal wall was not significantly different at 7.65 ± 3.95 nm/Pa. The displacement of the internal tracheal wall in the x direction (8.30 ± 2.26 nm/Pa) was also not significantly different from that in the z direction. Taken together these data suggest that mechanical coupling within the auditory organ is robust to different forcing regimes whether the majority of the force delivered to the system is sonic or generated by the mechanosensory neurons.

### Reconstruction of auditory organ motion

To fully reconstruct the relative motions of different parts of the auditory organ, we calculate the amplitudes of their displacement in response to sound and the phase of this motion relative to each other. We are mainly interested in the low SPL regime (40 dB SPL), since here the active amplification mechanism provides most of the force driving displacement.

As mentioned before, the anterior tympanal membrane and internal tracheal wall move almost exactly in anti-phase and there is relatively low variance in these values (Fig. 3D: angular mean ± angular SD: 168° ± 25°, N = 6). The relative phase between the x and the z direction of the tracheal wall shows a bimodal pattern, increasing the variance, but the mean suggests a smaller phase lag (Fig. 3D: angular mean ± angular SD: 138° ± 94°, N = 6). The mean phase differences suggest that on average the anterior tympanal membrane and the tracheal wall move in the opposite direction from each other in the z direction and that the internal tracheal wall traces an elliptical path internally within the organ (Fig. 4C).

**Figure 4.**
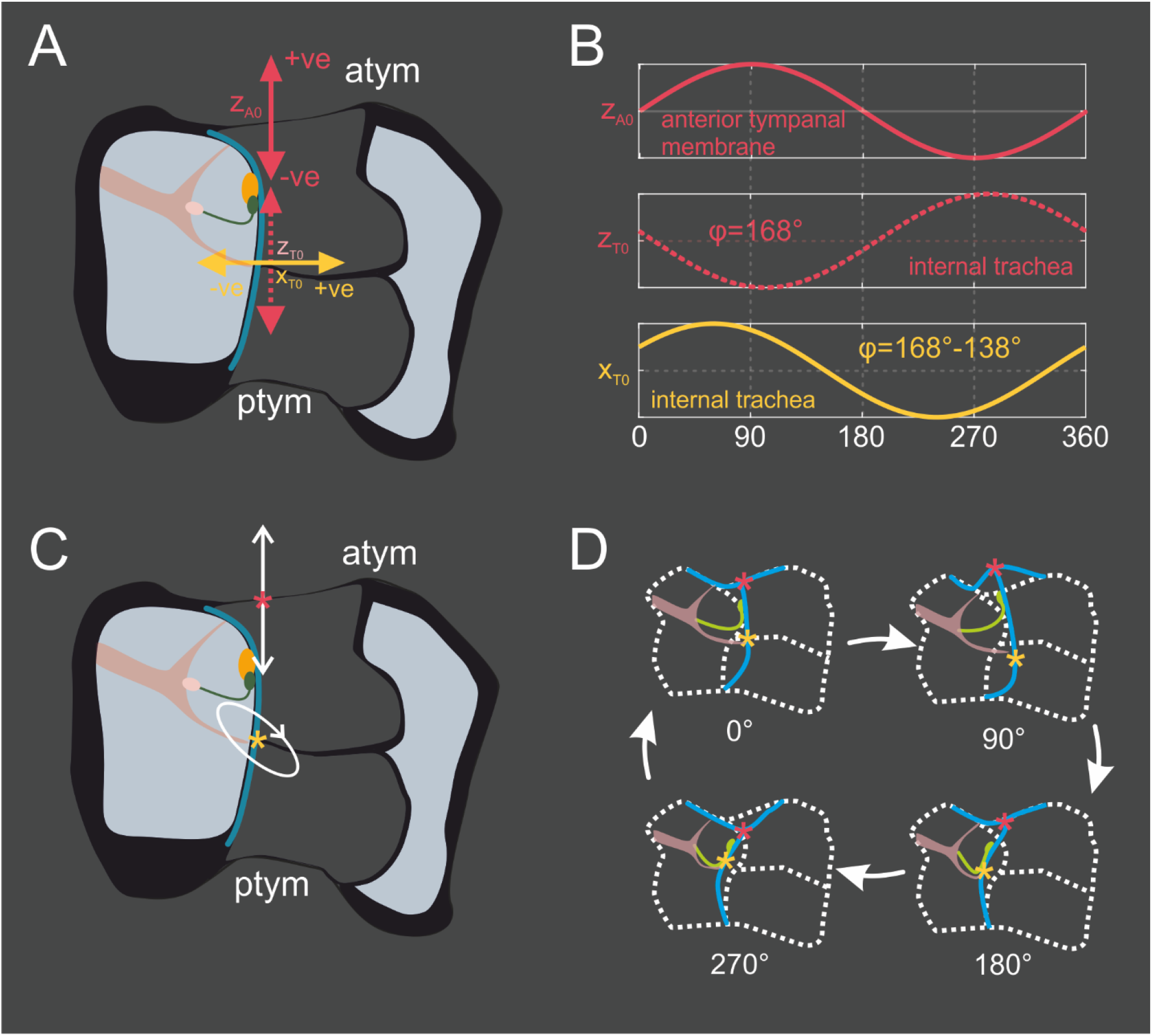
Reconstructing the motion of the tree cricket auditory organ at 40 dB SPL stimulation at the amplified frequency suggests bending motion of the dendritic cilia. (A) The OCT has enabled three measurements within the auditory organ. The measurements, their directions and 0 positions are depicted with respect to the anatomy here. In the z direction upwards is in the positive direction and vice versa. In the x direction, right is the positive direction and vice versa. (B) Here we plot one cycle of displacement we expect to observe at each of these points based on their relative phases. The external tympanal membrane and the internal tracheal wall are nearly anti-phase in the z direction and the motion of the internal tracheal wall in the x direction is more complicated. (C) If we consider its relative magnitude and phase in terms of motion within the x-z plane, the internal attachment point traces an elliptical path. (D) Based on this we can reconstruct the deformations that the auditory organ should undergo during a single cycle at the amplified frequency. At this stimulation level and frequency, the majority of the force (∼85%) moving the auditory organ is generated internally, by the action of molecular motors within mechanosensory neurons. At 0° both membranes are at resting position in the z direction but the internal membrane is deflected to the right. At 90°, the external membranes deflect upwards, and the internal membrane moves downwards. At this phase the two membranes are the furthest apart in the z direction. In the x-direction, the internal membrane remains in the right quadrant. At 180°, the two membranes move towards each other and arrive near their resting positions in the z direction, and the internal membrane has moved towards the left in the x direction. Finally at 270°, the two membranes are now the closest they will be to each other in the z direction and in the x direction, while the internal membrane remains within the left quadrant. Given that the dendritic cilia of the mechanosensory neurons (in green) are positioned at an angle to the tracheal wall and parallel to the membrane, we expect this motion to be caused by the cilium generating both vertical and lateral forces. The lateral motion alone could potentially be generated by a length change in the cilium, however coupled with the vertical motion as it is here, we believe at least some bending motion would be required.

In figure 4D, using these data, we reconstruct the motion within the auditory organ at the four phases (0°, 90°, 180° and 270°, figure 4D) of one cycle at the amplified frequency. Here, we see that the dendritic cilia, which drive active amplification, generate both lateral and vertical displacements within the organ. The cilia are expected to be the most mechanically compliant part of the neurons, given their relatively thin cross section (Moran et al., 1977) but they are also the force generating elements (Karak et al., 2015). Therefore, they must be generating both vertical and lateral forces. If the displacements observed were only lateral, one could posit that the displacements were generated solely by a change in their effective length as suggested in *Drosophila* (Hehlert et al., 2020). However, when the lateral motion is coupled with the vertical motion, this suggests that the dendritic cilia trace an elliptical path, and we have to posit that the cilium would also have to undergo bending motion to generate this displacement (Fig. 4D).

It is worth noting that the shape of this motion in an intact and alive auditory system will be quite similar no matter the SPL, just with different absolute amplitudes. The ratios of the amplitudes of the atym and tracheal wall motion, and their phases relative to each other do not change a great deal with stimulus SPL (Fig. 3 A-C). There are a few possible explanations for this, which are not mutually exclusive. This motion may be this mechanical system’s preferred motion (a mode, for instance) or the mechanical system may have developed so that both sonic and ciliary driving forces are delivered in the exact orientation so that they drive the tracheal wall in an elliptical path. While we cannot tease apart these possibilities, we can, however, say that the dendritic cilia do bend during active amplification.

We can also compare this vibrational behaviour to that observed *post mortem*, when the leg has been severed from the body. In a dead auditory organ, the amplitude of motion decreases significantly (Fig. S1). In addition, there is a large change in the phase difference between the z and x direction displacement of the tracheal wall. In a dead organ, the tracheal wall traces a significantly smaller elliptical path, which points in a different direction compared to the elliptical path observed in the live animal. Some of this change in vibrational behaviour is likely attributable to changes in impedance arising from changes in the severed acoustic trachea. It is also likely that the structural mechanics of the tympanal organ including the mechanosensory neurons is disrupted *post-mortem*. Our data suggests that in an alive animal, the tympanal organ generates a static and passive mechanical force due to its stiffness, in addition to a dynamic and active force due to action of motor molecules in the dendritic cilium. And that both these forces, together, generate an elliptical bending motion of the dendritic cilia.

### Motion during spontaneous oscillations

We were able to detect spontaneous oscillations from 5 crickets (Fig. S2). In two of these five, it was possible to make the full range of measurements required: record from single points on the anterior tympanal membrane in the z direction and from the internal tracheal in both the x and z directions (Fig. 5A). The noise floors for each of the measurements were variable but the peak amplitude displacement at the amplified frequency was similar in all three conditions, and to the behavior observed at low amplitude sound stimulation (Fig.3).

**Figure 5.**
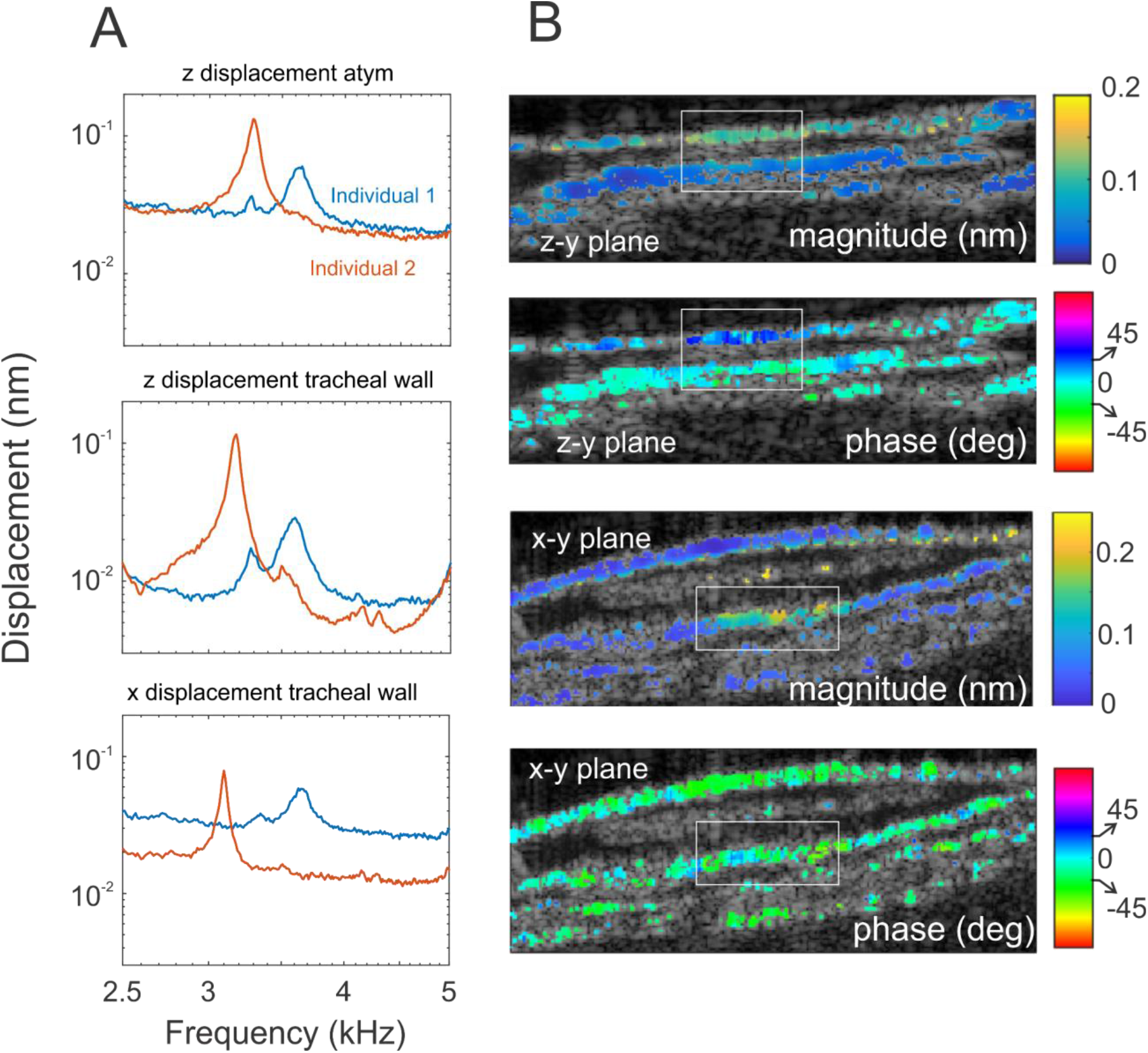
Spontaneous oscillations and transient evoked oscillations resemble sound driven responses. (A) Spontaneous oscillations were measurable in both the z and x directions in two animals. The amplitudes of the spontaneous displacements of the external anterior tympanal membrane (z direction) were similar to the amplitudes of the internal tracheal wall (in both the z and x directions). (B) We used a transient sound to entrain membrane motion. Subsequent oscillations were then sustained by the amplification mechanism and measured over a longer period (400 ms). Given the very brief drive, the behavior of this system will closely resemble spontaneous motion but the external drive provides a reference against which phase can be measured. M-scans show that the displacement magnitude and phase behavior of the organ resembles the driven behavior, suggesting that the reconstruction of organ motion described in the previous figure would also hold for spontaneous oscillations.

Reconstructing phase relationships during spontaneous oscillations was more difficult. To attempt to get a rough estimate of the phases of these three motions during spontaneous movements, we used a brief (2-10 ms) sound to entrain the system which we then measured over a much longer duration (400 ms). The initial stimulus can be used as a phase marker since we expect it to entrain oscillations initially for a brief duration. Using this phase marker, we measured 2D images of displacement phase (Fig. 5B) as before (Fig. 2D).

In this situation, displacement within the auditory organ is driven almost entirely by forces developed within the dendritic cilia of mechanosensory neurons. We expect that, because the entrainment is brief, the inherent phase incoherence of the active system eventually dominates. Nonetheless, even under these conditions, we find that the anterior tympanal membrane and the tracheal wall have a phase difference of ∼76°. This is a lower phase difference than that seen in the sound driven system, suggesting a smaller vertical displacement for this tympanal membrane during spontaneous oscillation. However, the important measurement is from the tracheal wall. Here, we observe a phase difference of ∼60° between its x and z displacement. This phase difference is also smaller, but still suggests that the tracheal wall undergoes a phase lagged lateral and vertical motion, thus tracing an elliptical path during spontaneous motion. Since the force for this path is generated by the dendritic cilia, it suggests that motion during purely spontaneous oscillations is also generated by the cilium bending. This evidence further shores up the suggestion that the dendritic cilia of tree cricket mechanosensory neurons bend as they produce forces for active amplification.

## Discussion

In the auditory organs of true crickets, mechanosensory neurons are not directly attached to external tympana or ‘eardrums’ (Yack, 2004; Yager, 1999). In this study, we have measured how vibrations travel from the tympanal membranes to the mechanosensory neurons within the auditory organ of a member of this group, the western tree-cricket (*Oecanthus californicus*). We found that while the mechanosensory neurons were not attached to the peripheral sound collecting structures (Fig. 1), they were mechanically well-coupled. Vibrations of the tympanal membranes travelled largely unattenuated to compliant internal structures such as the acoustic trachea (Fig. 3). Other stiffer structures such as the wall of the acoustic vesicle remained unmoved (Fig. 2). Since the mechanosensory neurons are suspended between the vibrating acoustic trachea and the immobile auditory vesicle wall, we can surmise that vibrations were transmitted to them (Fig. 4). We also found that the source of the vibrations (external source vs. the physiological amplifying action of mechanosensory neurons) did not have a significant effect on transmission within the alive organ (Fig. 3, 5, S1).

### Morphology and mechanics of related auditory organs

An important acoustically active sister group to the tree crickets are the Tettigoniids, commonly known as katydids or bushcrickets (Song et al., 2020). In katydids, the auditory organ is known as the crista acustica (Montealegre-Z. et al., 2012) and has a similar structure with one crucial difference, the mechanosensory neurons are not attached to the wall of the auditory vesicle (Strauß, 2019). In fact, phylogenetic evidence suggests that the two organs evolved independently from a precursor intermediate organ (Song et al., 2020; Strauß and Lakes-Harlan, 2014).

The mechanics of the Tettigoniid auditory organ have been proposed to be quite distinct and have some features that strongly resemble the auditory mechanics of the mammalian cochlea (Montealegre-Z. et al., 2012; Scherberich et al., 2020). In this system, it has been suggested that a part of the tympanal membrane acts as a lever, pushing on the fluid in the auditory vesicle. This mechanical action is then thought to generate a fluid wave which stops at different locations within the vesicle (Montealegre-Z. et al., 2012). The stopping location of each frequency is tonotopically arranged, thus each frequency excites a distinct set of mechanosensory neurons (Montealegre-Z. et al., 2012; Scherberich et al., 2020).

In this schema, the acoustic trachea within the crista acustica has little role to play in the mechanical behaviour of the organ (Montealegre-Z. et al., 2012). Indeed, until recently the function of trachea in cricket and katydid organs has been thought to be primarily acoustic in nature, i.e. they were thought to serve largely as tubes that carry sound to the back surface of tympana, acting as secondary sound inputs which are largely only relevant to the development of directionality in hearing (Celiker et al., 2020; Jonsson et al., 2016; Michelsen and Larsen, 2008, 1978). However, our data clearly demonstrates that cricket trachea do have a mechanical role. Using OCT based vibrometry we were able to measure the motions of the part of the acoustic trachea that is attached to the tympanal membranes. Our data shows that, within the auditory organ the internal tracheal wall moves at amplitudes similar to that of the tympana, and couples the motion of the tympana to that of the mechanosensory neurons. It is thus possible that the motion observed in the katydid organ (Montealegre-Z. et al., 2012; Scherberich et al., 2020) may also be a structural rather than a fluid wave or may contain both elements, which would be more similar to current models of the mammalian cochlea (Bowling and Meaud, 2018; Meaud and Grosh, 2012). We propose that OCT based depth resolved vibrometry would be a more sensitive technique than surface-level vibrometry to test the similarities and differences between these organs.

The other well-known feature of vertebrate hearing is active auditory amplification, and while the evidence for a travelling wave and tonotopy is quite strong in the Tettigoniid auditory organ, no clear evidence has emerged for active amplification (Mhatre, 2015). In contrast, active amplification has now been observed in *Oecanthus californicus* and two other tree cricket species, all of which fall within the Gryllidae lineage (Mhatre et al., 2016; Mhatre and Robert, 2013; Morley and Mason, 2015). Thus, one possibility is that two closely related cricket lineages possess two different features of mammalian hearing, which have evolved entirely independently, and another is that current methods do not have the sensitivity to detect amplification in Tettigoniids. We propose that the OCT measurement paradigm we present here would provide a more sensitive method to test whether both processes may be present simultaneously in both or either group. As now demonstrated in Tettigoniids, OCT is invaluable to measuring internal mechanics in detail (Scherberich et al., 2020, 2016; Vavakou et al., 2021) and will be crucial in teasing apart the different contributing mechanisms that might determine the center frequency and sharpness of auditory tuning.

### Active amplification in insect ears: organelles and their molecular biology

More interestingly, active auditory amplification in insects and mammals are based on entirely different organelle structures, and involve entirely different molecular players (Hehlert et al., 2020; Yack, 2004). The specific channels, motor proteins and structural mechanisms underlying active auditory amplification in mammals are still being investigated and are topics of great debate (Ashmore et al., 2010). However, one thing is well established, the kinocilium which is the only true cilium found on hair cells is crucial for patterning during development but is resorbed in adults and hence plays no part in auditory amplification (Eatock, 2006).

Data from *Drosophila* shows that in insects, however, active amplification is based on a true cilium; specifically on ciliated dendrites in auditory neurons (Gopfert et al., 2005) and the molecular motors that generate amplifying forces are the axonemal dyneins found in all cilia (Karak et al., 2015). In typical motile cilia, in sperm or swimming algae for instance, axonemal dynein motors cause parallel microtubule pairs to slide against each other. The microtubules sliding past each other, then cause the ciliary shaft to bend (Lin and Nicastro, 2018; Satir et al., 2014). The timing of sliding is thought to be structurally coordinated, so that it occurs alternately on opposite sides of the cilium, making cilia bend towards each side in alternation, and giving rise to a beating motion which has a specific frequency (Satir et al., 2014; Yagi and Kamiya, 1995). The mechanism by which the beat frequency is determined has been studied extensively (Satir et al., 2014), and the emerging consensus is that this mechanism shares several of the non-linear dynamical features observed in amplified hearing (Klindt et al., 2016; Ma et al., 2014). An early hypothesis for auditory amplification was, in fact, based on this bending and beating activity of the kinocilia which are present in amphibian ears (Camalet et al., 2000).

An important question that then arises is, is this ciliary beating responsible for dendritic cilia-based amplification in insect auditory organs? Data from some insect systems does suggest that dendritic cilia undergo a bending motion (Hummel et al., 2016; Moran et al., 1977), while other models favour a motion along the long axis of the dendrite (Hehlert et al., 2020). It is difficult to disentangle between the two models since, in insect auditory organs, the dendritic cilia of mechanosensory neurons are not unattached as they are in hair cells or sperm or swimming cilia. They are instead mechanically coupled to accessory cells (Yack, 2004), which could alter the shape of their motion during transduction and amplification.

Understanding this motion is crucial, however, as it would constrain models of force transmission and ion channel gating in the insect auditory system (Effertz et al., 2012; Hummel et al., 2016). In the gryllid auditory organ, mechanosensory neurons are arranged in a sheet-like structure that runs along the axis of the foreleg, parallel to the two tympanal membranes (Michel, 1974). In such a structure, in the tree cricket, it was possible to distinguish between bending and single axis movements. We observed significant neuronal movement along at least two axes, with the base of the neuron undergoing an elliptical movement. Such elliptical motion is observed both when the mechanical forces are generated by external sounds and when they are generated by the internal amplification process. This motion reduces in amplitude and changes phase *post-mortem*. This data together suggests that at least in the tree cricket, the dendritic cilia do undergo a bending and beating motion. Similar bending of mechanosensory neurons has been observed in Tettigoniids as well, where the neurons are arranged in a similar row like fashion (Hummel et al., 2016). If indeed, fly neurons behave differently than cricket neurons, this suggests a greater diversity of force transmission and ion channel gating mechanisms than previously appreciated even within the insects.

This finding, however, does lead to some exciting conclusions about the evolution of active amplification. The cilium is an ancient and phylogenetically widely distributed organelle. Ciliary bending and beating motility plays a crucial functional role in an incredible diversity of contexts: swimming, embryonic symmetry breaking, cerebrospinal fluid flow and even in olfaction (Ringers et al., 2020; Wan and Jékely, 2020). Our measurements here suggest that insect ears can exploit the same ancient mechanism for amplification even in an auditory context. While much remains to be understood about how this behavior functions in the huge diversity of insect auditory organs, it is nonetheless intriguing to consider that the evolutionary origin of auditory amplification may well precede the evolution of audition altogether.

## Methods

### Animals

Vibrometric data were collected from 6 wild-caught *Oecanthus californicus* individuals (5 males, 1 female). Animals were anaesthetised using CO_2_ and mounted on a single 3D printed plastic block using wax as previously described (Mhatre et al., 2016; Mhatre and Robert, 2013) and the auditory organ on the foreleg tibia was placed in the path of the laser.

### Volumetric OCT and vibrometry (VOCTV)

The VOCTV method which has been previously described and is a custom built swept-source optical coherence tomography system (Lee et al., 2015) that allowed us to measure vibrations throughout the volume of the tree cricket’s tibial auditory organ. Unlike standard laser Doppler vibrometry (LDV), OCT allows for depth resolved vibration measurement within tissue. VOCTV like LDV is based on an interferometric measurement, but it uses long wavelength light (1300 nm) which penetrates tissue (Gao et al., 2012; Lee et al., 2015). Additionally OCT generally, and VOCTV specifically uses broad-band low-coherence light which allows us to use coherence length to reject reflections from all but a thin section of tissue (Drexler et al., 2014; Lee et al., 2015). This in turn allows us to measure vibrations even from subsurface structures. The imaging resolution of the system is ∼9.8 and 12.5 μm in the lateral and axial dimensions respectively (FWHM, refractive index = 1.4) and the temporal sampling rate 100 kHz (Dewey et al., 2021).

Data from the VOCTV system was stored in custom format, and analyzed and plotted using custom Matlab scripts. Data from VOCTV contains two basic components, the reflectivity of the tissue at different depths along the laser beam path, and the vibrational component, i.e. displacement of the tissue at these positions, which is calculated from interleaved scans. Two and three dimensional versions of these data can be acquired by sweeping the laser beam along the tissue and by spatially reconstructing both reflectivity and vibrational patterns.

### 3D reconstruction

For 3D reconstruction, vibrometric data was omitted and only the data which reflects the amplitude of the reflection from different depths with the tissue in the scanned volume was imported into the software Slicer 4.10.1. This data was threshold segmented by hand and was converted into a STL file. This file was imported into Meshlab v2016.12 and where the mesh was repaired and smoothed and prepared for final display.

### Sound stimulation paradigms

Sound stimuli were produced using a Fostex speaker (FT17H) and calibrated in the free field situation just above the foreleg of the insect in the preparation before each experiment using a Brüel & Kjær 1/8” microphone (Type 4138). For the single point measurements, we measured a 1.5 kHz span around the amplified frequency with logarithmically increasing frequencies. The sound stimulus was a ramped 100 ms pure tone and the OCT measurement was run for the same duration. Each measurement was made at least 5 times and the responses averaged.

For the 2D measurements, the stimulus and measurement duration remained the same, however, no averaging was carried out. In these scans, displacement data was filtered in based on the amplitude of the reflection from the tissue. After testing a few settings, we display only those displacement data that were measured from parts of the tissue that were at least 58% as reflective as the highest reflection level measured.

To make measurements in two planes (Fig. 2) the preparation was rotated by hand using the animals atym and ptym structures to guide the rotation. It is possible that the two planes are not exactly orthogonal to each other. However, given the small dimensions of the whole auditory organ (1-2 mm), large changes in angle would be noticeable since they would exit the organ in very different planes. Given the similarity of the 2D scan patterns, it is likely that the measurements were relatively similar in angle.

While measuring spontaneous oscillations, no sound stimulus was produced, and a vibrometric measurement was made for 100 ms, the spectrum for each measurement was calculated and then averages of 400 measurements were made. For the 2D scans made for the transient entrainment experiment, the stimulation duration was 2 ms (xz plane) and 10 ms (xy plane) at 100 dB SPL. No averaging was carried out.

## Acknowledgements

We would like to thank Christopher Bergevin and the Mechanics of Hearing community for fostering this collaboration. The contributions of Frank Macias-Escriva, Sangmin Kim and Wihan Kim to the OCT software and training of NM are also gratefully acknowledged.

We would also like to thank the following funders for their support of this research: NSERC Discovery (Grant no. : 687216) and an NSERC Canada research chair (Grant no. : 693206) to NM; an NIH/NIDCD F32 fellowship (DC016211) to JBD; a NSERC Discovery (Grant no. : 2018-04917) to AM; NIH/NIDCD Grants (R01 DC014450, R01 DC013774, and R01 DC017741) to JSO and an NIH/NIBIB grant (R01 EB027113) to BEA.

## Figures

**Supplementary figure 1:**
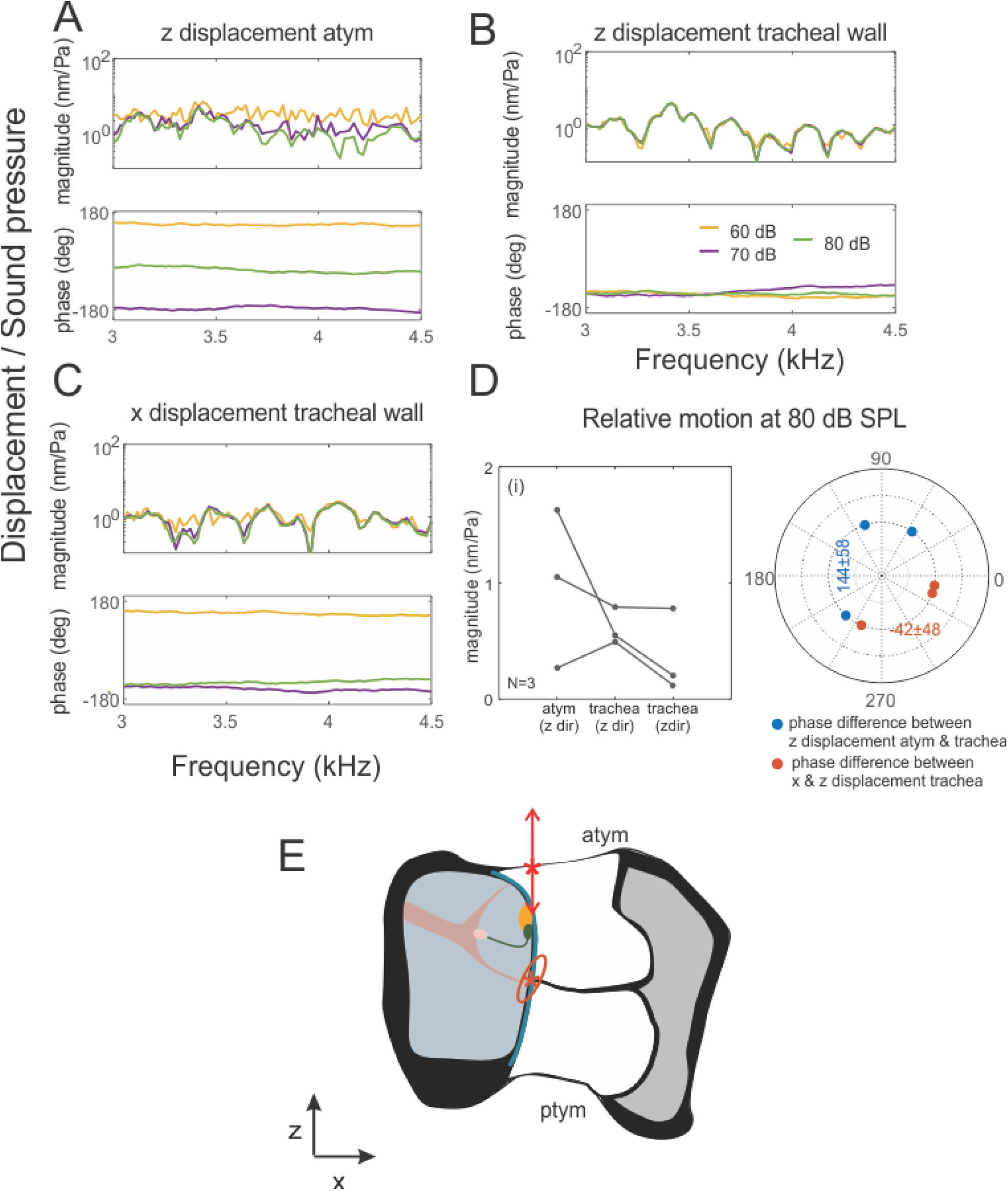
The frequency response of the tree cricket auditory organ becomes linear *post mortem*. We investigated whether the non-linearity of the tree cricket auditory organ was dependent upon its physiology by severing the leg from the body and measuring sensitivity from the same positions in the organ as in figure 3; from the point of maximal displacement on the (A) external anterior tympanal membrane, and (B, C) the internal tracheal wall (depicted by *). The sensitivity or displacement relative to sound pressure level was measured at these points at different stimulation levels, along the (A, B) z and (C) the x axis. When the leg was severed from the body we saw a strong decline in levels of motion even at 80 dB SPL (atym z direction (dead): 0.98 ± 0.68 nm/Pa vs, the same individuals when alive: 10.14 ±5.71 nm/Pa (N=3, paired 1 tailed t-test, P=0.061); tracheal wall z direction (dead): 0.61 ± 0.16 nm/Pa vs. (alive): 8.7 ± 4.68 nm/Pa (N=3, paired 1 tailed t-test, P=0.045); tracheal wall x direction (dead): 0.37±0.37 nm/Pa vs. (alive): 9.57 ± 0.83 nm/Pa (N=3, paired 1 tailed t-test, P=0.001)). Where measurements were above the noise levels, we saw no evidence of compressive non-linearity and level dependent tuning in all three measurements including at the attachment point of the tympanal organ. (D) We measured (i) the magnitude of the motion of the two points in the two direction at a stimulation level of 80 dB SPL, from 3 individuals near conspecific frequency and (ii) their phases relative to each other. Using the average magnitude and phase difference of displacement at the two points we were able to (E) reconstruct the shape of motion of the two parts of the auditory organ during sound stimulation. The external atymp and internal tracheal wall still moved out of phase with each other, however, the phase lag was reduced to 114 ± 58°, as was the phase lag between the x and z direction of the internal tracheal wall. In addition, the shape of the motion of the internal tracheal wall also changed, the elliptical path in the dead leg was pointed in the opposite direction.

**Supplementary figure 2:**
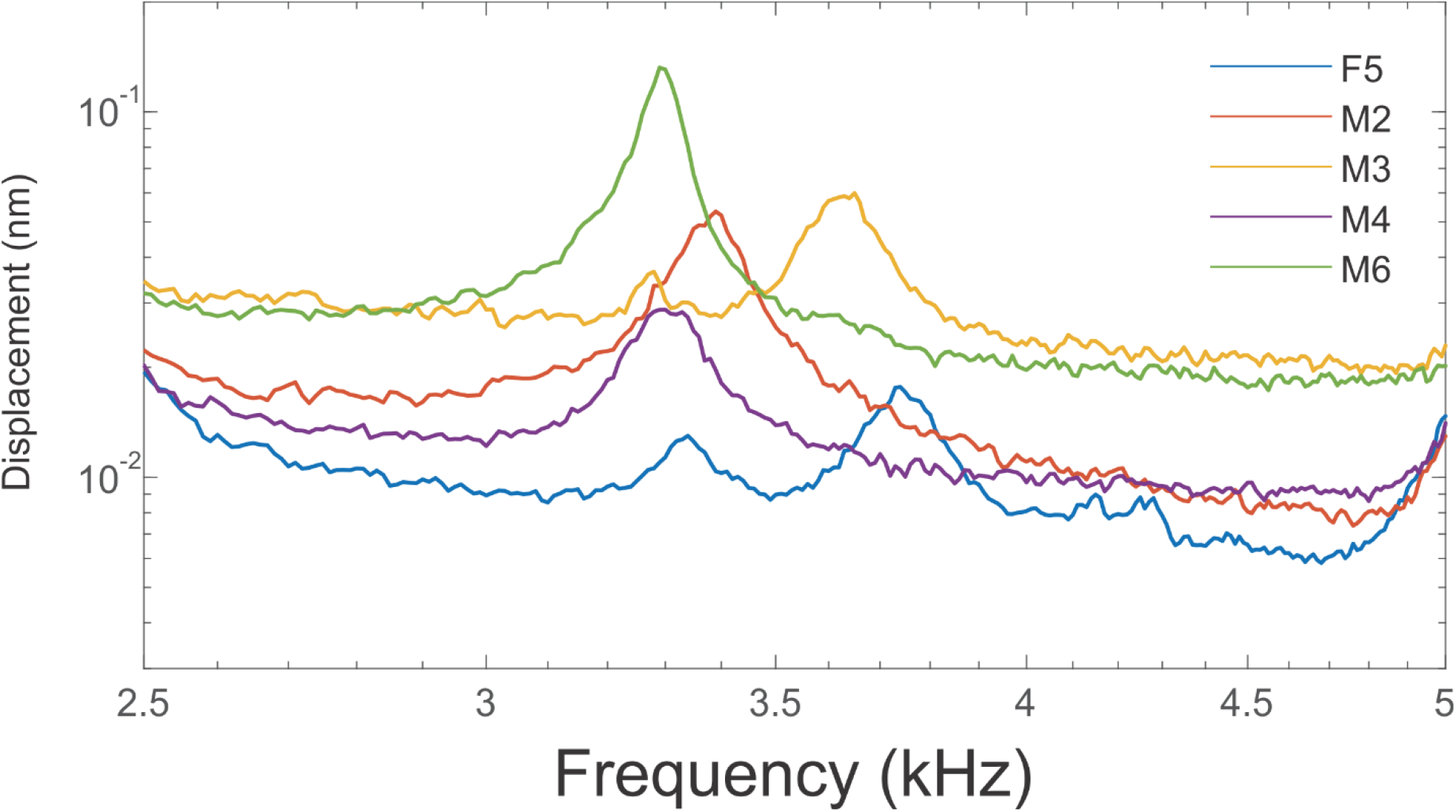
Spontaneous oscillations from the anterior tympanal membrane. Spontaneous oscillations in the z direction were tested for in all individuals from the anterior tympanal membranes and observed in 5 out of the 6 individuals we tested. Measurements from this membrane were usually the first measurements made from an experimental animal. As the experimental procedure progressed, we observed a gradual decline in animal health, and sound driven measurements were prioritized. Thus, we were able to make all three measurements (external membrane z axis & internal trachea z and x axis) from only 2 animals (Fig. 5).

